# An exosome-based gene delivery platform for cell-specific CRISPR/Cas9 genome editing

**DOI:** 10.1101/2023.06.09.542202

**Authors:** Sunil Dubey, Zhe Chen, Austin Talis, Andrei Molotkov, Alessandra Ali, Akiva Mintz, Fatemeh Momen-Heravi

## Abstract

Exosomes are naturally occurring vesicles that have the potential to be manipulated to become promising drug delivery vehicles for on-demand in vitro and in vivo gene editing. Here, we developed the modular safeEXO platform, a prototype exosome delivery vehicle that is mostly devoid of endogenous RNA and can efficaciously deliver RNA and ribonucleoprotein (RNP) complexes to their intended intracellular targets manifested by downstream biologic activity. We also successfully engineered producer cells to produce safeEXO vehicles that contain endogenous Cas9 (safeEXO-CAS) to effectively deliver efficient ribonucleoprotein (RNP)-mediated CRISPR genome editing machinery to organs or diseased cells in vitro and in vivo. We confirmed that safeEXO-CAS exosomes could co-deliver ssDNA, sgRNA and siRNA, and efficaciously mediate gene insertion in a dose-dependent manner. We demonstrated the potential to target safeEXO-CAS exosomes by engineering exosomes to express a tissue-specific moiety, integrin alpha-6 (safeEXO-CAS-ITGA6), which increased their uptake to lung epithelial cells in vitro and in vivo. We tested the ability of safeEXO-CAS-ITGA6 loaded with EMX1 sgRNAs to induce lung-targeted editing in mice, which demonstrated significant gene editing in the lungs with no signs of morbidity or detectable changes in immune cell populations. Our results demonstrate that our modular safeEXO platform represents a targetable, safe and efficacious vehicle to deliver nucleic acid-based therapeutics that successfully reach their intracellular targets. Furthermore, safeEXO producer cells can be genetically manipulated to produce safeEXO vehicles containing CRISPR machinery for more efficient RNP-mediated genome editing. This platform has the potential to improve current therapies and increase the landscape of treatment for various human diseases using RNAi and CRISPR approaches.

## INTRODUCTION

Exosomes are nanosized (70-200 nm) membrane-bound vesicles found in biofluids and secreted by nearly all types of cells in the cellular microenvironment ^1-3^. They naturally carry biomacromolecules—including different RNAs (mRNAs, regulatory microRNAs (miRNAs)), DNAs, lipids, and proteins—and can efficiently deliver their cargos to recipient cells, mediating cellular communication and functionality ^4^. Previous work by our group and others has shown the advantages of using exosomes for drug delivery. Exosomes 1) are small and have a high efficiency for delivery due to their similarity to cell membranes; 2) are biocompatible, non-immunogenic, and non-toxic, even with repeated in vivo injections ^5^; 3) are stable even after several freeze and thaw cycles; 4) contain a lipid bilayer that protects their protein and RNA cargos from enzymes such as proteases and RNases ^2^; 5) have a slightly negative zeta potential, leading to long circulation ^6^; and 6) exhibit an increased capacity to escape degradation or clearance by the immune system ^7,8^. Thus, exogenous exosomes are being developed for their potential to deliver RNA interference (RNAi), miRNAs, and mRNA by our group and others, ^5,9,10^ and hold great potential for future therapeutic uses.

The CRISPR/Cas genome editing system has been utilized extensively in recent years for its ability to produce targeted genome editing. The technology is rapidly maturing as a clinical grade technology for select genetic diseases and is actively being explored for a myriad of diseases. However, challenges remain in its widespread therapeutic use, partly due to the lack of a targeted, reliable, and safe delivery method for the requisite gene editing components. Exosomes present a promising delivery vehicle for genome editing cargo, but innovative approaches are needed to overcome several challenges in order to successfully utilize them as a scalable, targeted, and reliable vehicle for CRISPR/Cas machinery to efficiently facilitate gene editing without unwanted biological effects or uncharacterized endogenous cargo.

CRISPR-based editing induces double-stranded DNA (dsDNA) breaks in a targeted fashion, prompting resultant DNA repair via either nonhomologous end joining (NHEJ) or homology directed repair (HDR). NHEJ occurs without a template and thus may be exploited for gene knockout by deletion, frameshift or the introduction of new STOP-codons via mutation ^11,12^. NHEJ requires a minimal Cas endonuclease and single guide RNA (sgRNA). Cas plus a single sgRNA are adequate for gene disruption/mutation, while Cas plus two sgRNAs are needed for gene deletion. HDR, which inserts new genetic material, requires Cas, sgRNAs along with a ‘spare’ DNA template for targeted genetic insertion. HDR thus provides an opportunity to knock-in specific DNA sequences at targeted locations in the genome to correct genetic diseases or introduce new functional proteins, although it is much less efficient compared to NHEJ ^13,14^.

One of the main challenges of using CRISPR/Cas for the treatment of human diseases is the lack of efficient delivery methods for the requisite complex multicomponent machinery that can target specific cells and organs in vivo and avoid gene edits in non-target cells or genetic loci ^15^. CRISPR/Cas can be delivered locally or systemically via gene-based delivery (DNA plasmids or viruses encoding Cas and sgRNAs), RNA-based delivery (mRNA encoding Cas along with a synthetic sgRNA), or formed Ribonucleoprotein (RNP) complexes consisting of Cas protein prebound to synthetic sgRNA. Of these three methods, RNP-based delivery appears to be superior and most specific to editing the targeted genome site^16^ because it does not rely on uncontrolled CAS integration or expression, although it is also the most difficult to implement as a therapeutic due to the complexities of manufacturing and efficiently delivering intact, functional recombinant Cas endonuclease proteins together with the requisite sgRNA and DNA templates specifically to the site of disease.

Although both viral and non-viral approaches have been adopted for in vivo delivery of CRISPR/Cas machinery, the effective in vivo delivery of multiple CRISPR components into host cells remains a major challenge. Adenovirus (AV) is an efficient transducing agent used for CRISPR/Cas-mediated genome editing, however, this method can elicit a significant immune response in the host ^17^. Lentiviral vectors are widely used for therapeutic delivery, though their integration into the genome makes them suboptimal for gene editing purposes as long-lasting expression of Cas protein and sgRNA is considered to be unfavorable for the on-target/off-target ratio of indel formation ^18^. Further, viral vectors are limited in terms of cargo carrying capacity and cell tropism. These shortcomings present difficulties regarding the distribution and dosage of genome editing nucleases in vivo, leading to off-target mutation profiles that may be difficult to predict ^18,19^. Non-viral synthetic vectors are another class of delivery vehicles which lack tissue tropism, yet they may provide targeted cell/organ-specific delivery if complexed with targeting moieties such as peptides or antibodies ^20^. Precise targeting, however, is particularly difficult to achieve, as incorporation of additional biomolecules to a delivery vector alongside the CRISPR components increases packaging complexity ^21^. Disadvantages of synthetic delivery vectors include issues with biocompatibility and toxicity, immunogenic potential, and problems with therapeutic cargo release ^19^. Thus, current implementations of CRISPR/Cas machinery via viral- and non-viral delivery present challenges that prevent the full therapeutic exploitation of gene editing and novel solutions are needed to translate the scientific progress in gene editing to benefit patients.

In this work, we engineered exosomes for effective, targeted, and scalable delivery of CRISPR/Cas machinery and demonstrate efficient gene editing in vitro and in vivo. We generated a novel exosome-based drug delivery platform **(Fig. 1)** engineered (1) to minimize endogenous nucleic acid cargo (safeEXO); (2) to endogenously carry an active Cas9 protein (safeEXO-CAS); and (3) to express tissue specific targeting moieties. Our safeEXO-CAS loaded exosomes efficiently edited recipient cells in vitro and in vivo using NHEJ-mediated disruption or HDR-mediated insertions without inducing immunogenic response or off-target effects.

**Fig 1.**
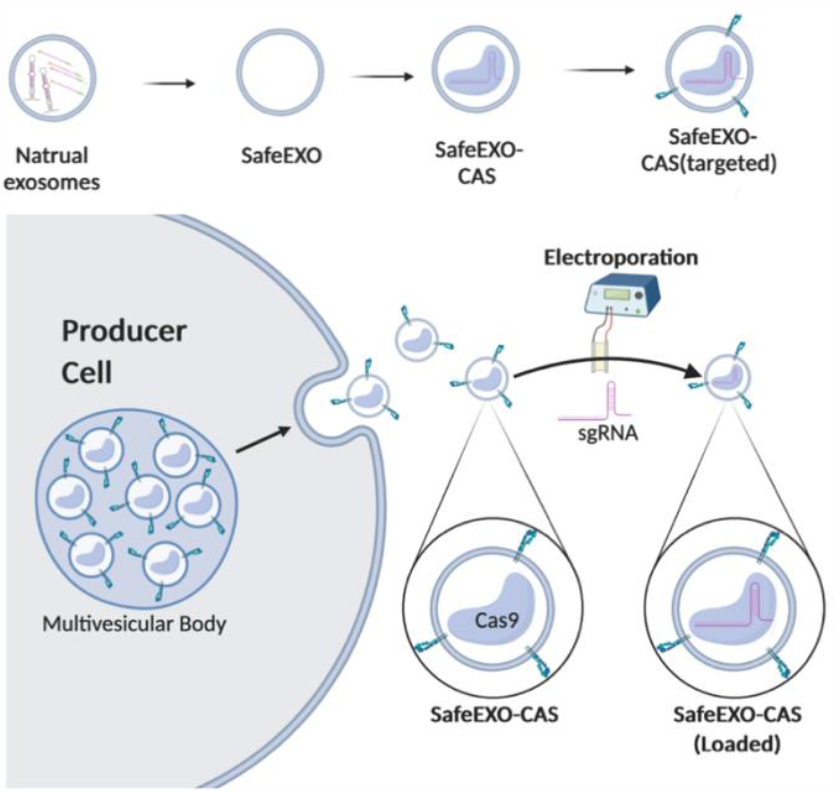
Schematic representation of safeEXO-CAS exosomes.

## RESULTS

### SafeEXO – Creating exosomes devoid of endogenous RNA

It is well documented that exosomes, based on their cellular origin, carry specific and considerable amounts of nucleic acids such as mRNA and miRs ^23-25^. RNA carried by exosomes have been implicated in propagating many diseases, including cancer and infectious disease. Thus, the presence of significant endogenous biologically active RNA cargo introduces complications in the manufacturing and characterization of exosomes and might introduce potential adverse effects that complicate their use as therapeutic vehicles. Here, we aimed to overcome this limitation by producing exosomes which are almost devoid of endogenous nucleic acid cargo that can be used for the loading and delivery of only the intended miRs, mRNA, and sgRNA being used for therapeutic purposes. To accomplish this in a scalable manner, we knocked out components of the endogenous exosome RNA loading machinery in exosome producer cells. Heterogeneous nuclear ribonucleoprotein (hnRNPA2B1), binds to RNA through the recognition of specific motifs, controlling its loading into exosomes ^26^. Drosha plays a role in miR processing and loading into exosomes ^27^. ALIX plays a role in the loading of exosomes with miRs and mRNA^28^. Silencing hnRNPA2B1, ALIX, or Drosha resulted in a significant reduction of endogenous RNA in the exosomes and production of exosomes practically devoid of endogenous nucleic acids **(Fig.2a)**. Elimination of hnRNPA2B1, ALIX, and Drosha in producer cells did not change the number or quality of exosomes produced. Exosomes isolated from hnRNPA2B1, ALIX, and DROSHA KO cell lines demonstrated normal protein/particle ratio **(Fig.2b)**, and no significant difference between levels of exosome enriched tetraspanins CD63, and CD81 via flow cytometry (**Fig.2c, d)** compared to parental cell exosomes. While producer cells with ALIX KO showed normal growth patterns, cells with Drosha KO and hnRNPA2B1 KO exhibited less proliferation **(Supplementary Fig.1)**. Thus, exosomes produced from ALIX-KO cells, referred herein as safeEXO, that are nearly devoid of potentially unwanted endogenous RNA, were used for further experiments and as the base vehicle for delivery of RNA and genome editing machinery. SafeEXO exosome production was accomplished in 3 independent cell lines, demonstrating the broad applicability generating exosomes mostly devoid of RNA, including from standard exosome producer cells or cancer cells (**Supplementary Fig.2**). The biological significance of the safeEXO platform was tested by exposing cancer cell-derived normal or safeEXO exosomes to CAL27 cells, which have been shown to increase proliferation when exposed to cancer produced exosomes. safeEXO exosomes generated from CAL27 did not increase proliferation of CAL27 cancer cells, in contrast to the natural exosomes that were produced by the same background cell line (**Supplementary Fig.3**). To confirm that safeEXO vehicles maintained their ability to deliver exogenously loaded RNA into cells, we exposed cells to safeEXO exosomes loaded with cel-miR-39 and demonstrated robust cell uptake and intracellular targeting manifested by downstream functionality **(Supplementary Fig.4)**.

**Fig 2.**
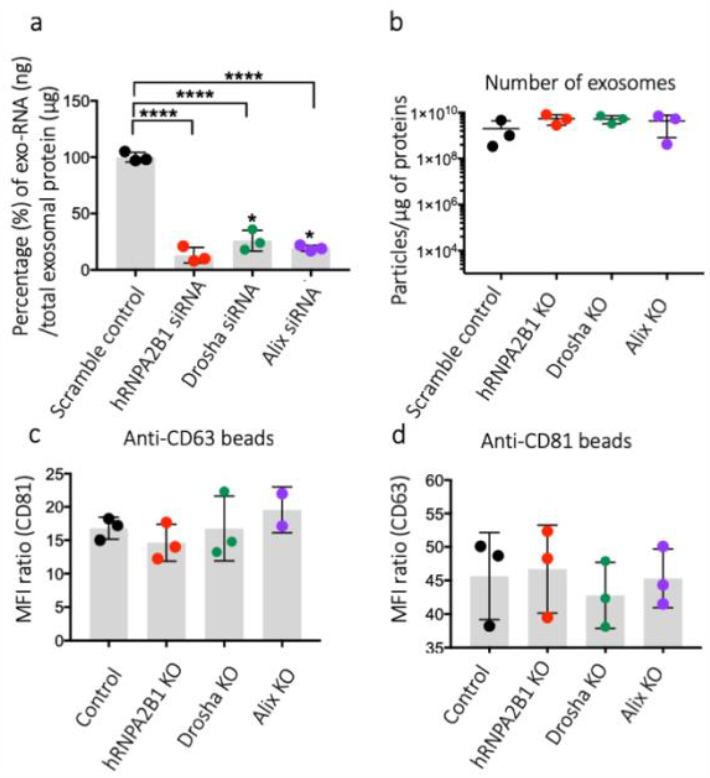
SafeEXO-CAS characterization and preparation. **a** Percentage change of total exo-RNA (ng) normalized based on the exosomal protein in exosomes isolated from siRNA silenced hnRNPA2B1, ALIX, and Drosha THP-1 monocytes. **b** Number of exosomes produced per μg of exosome protein from hnRNPA2B1, ALIX, and Drosha knockout THP-1 monocytes. **c, e** Flow cytometric analysis of exosomes from hnRNPA2B1, ALIX, and Drosha knockout for the presence of exosome enriched markers CD63 and CD81. SafeEXO-CAS exosomes were labeled with anti-CD81 or anti-CD63 magnetic beads and the levels of CD81 and CD63 were quantified by flow cytometry.

### SafeEXO vehicles engineered to endogenously express Cas9

Genome editing represents a major interest for basic and translational studies. To generate Cas9 safeEXO vehicles, ALIX-KO safeEXO producer cells were further engineered to knock-in Cas9 and used to generate RNA-cleared, exosomes containing endogenous Cas9 (safeEXO-CAS). exosomes that express canonical exosomal markers (CD63 and TSG101) and CAS9 **(Fig.3a)**. SafeEXO-CAS exosomes showed a mean size of around 170 nm measured by Nanosight analysis **(Fig.3b)**. To confirm cellular targeting and internalization by target cells, safeEXO-CAS vehicles were labeled with PKH26 or Di-8-ANEPPS. PKH26 constitutively fluoresces but Di-8-ANEPPS conditionally fluoresces upon internalization into the target cell,^29^ thus allowing the target uptake of fluorescently labeled safeEXO-CAS exosomes to be visualized and quantified via microscopy and flow cytometry. After a 1h co-culture, safeEXO-CAS exosomes were taken up efficiently by the recipient cells **(Fig. 3c)**. Flow cytometry confirmed a dose-dependent increase in the safeEXO-CAS uptake after co-culture **(Fig.3d)**.

**Fig 3.**
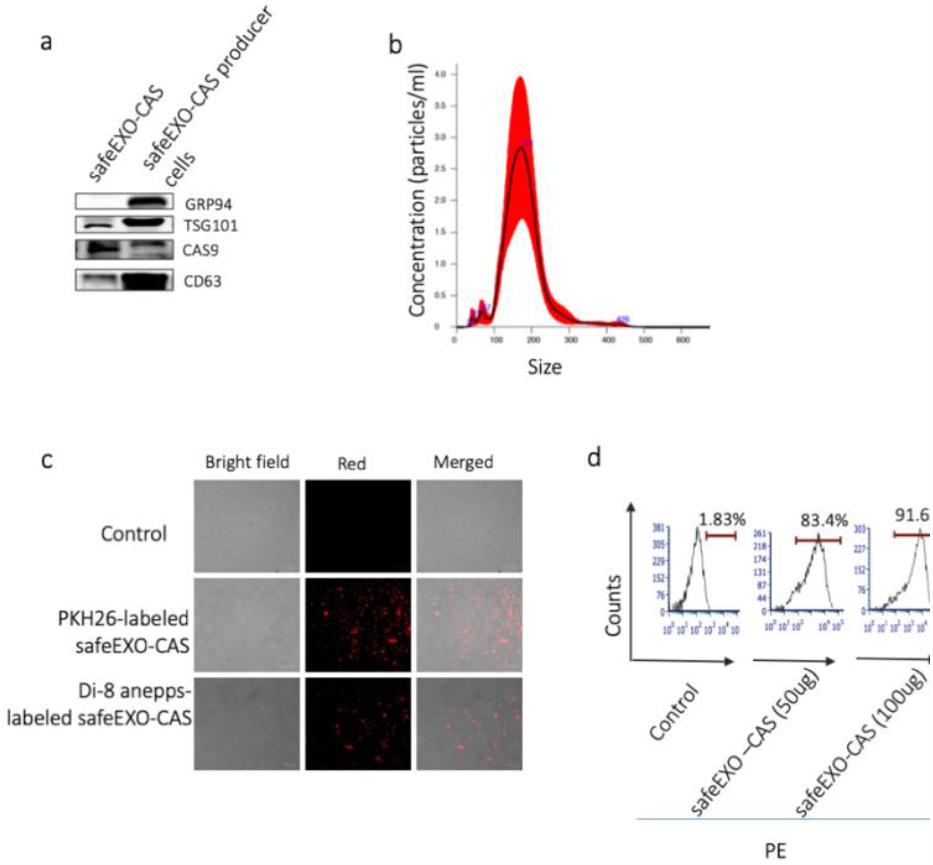
SafeEXO-CAS characterization and uptake. **a** Western blot analysis of exosome markers CD63 and TSG101, along with CAS9 and GRP84 (negative control) in safeEXO-CAS exosomes from THP-1 monocyte producer cells with and without ALIX knockout. **b** Size of safeEXO-CAS exosomes quantified by NanoSight. **c** Fluorescence microscopy of THP-1 cell line co-cultured with PKH26 and Di-8-ANEPPS labeled safeEXO-CAS after 1 hour. **e** Flow cytometry analysis of Di-8-ANEPPS labeled safeEXO-CAS uptake in THP-1 cell line after 1h co-culture. Experiments were repeated twice and data are represented as mean ± standard deviation (SD). \*\*\*\**p* ≤ 0.0001, One-way ANOVA was use for comparison of multiple group and Student’s T test was used for pairwise comparison.

### SafeEXO-CAS vehicles mediate efficient CRISPR-based NHEJ genomic editing

Next, safeEXO-CAS exosomes were tested for their potential to efficiently induce NHEJ genome editing using a GFP HEK293T reporter cell line that was exposed to safeEXO-CAS exosomes loaded with sgRNA directed at disrupting GFP. Genome editing was demonstrated by a significant decrease in GFP expression in target cells as demonstrated by fluorescent microscopy and flow-cytometry **(Fig. 4a, 4b)**. Indel induction in the target cell line by T7 endonuclease assay **(Supplementary Figure 5)**.

**Fig 4.**
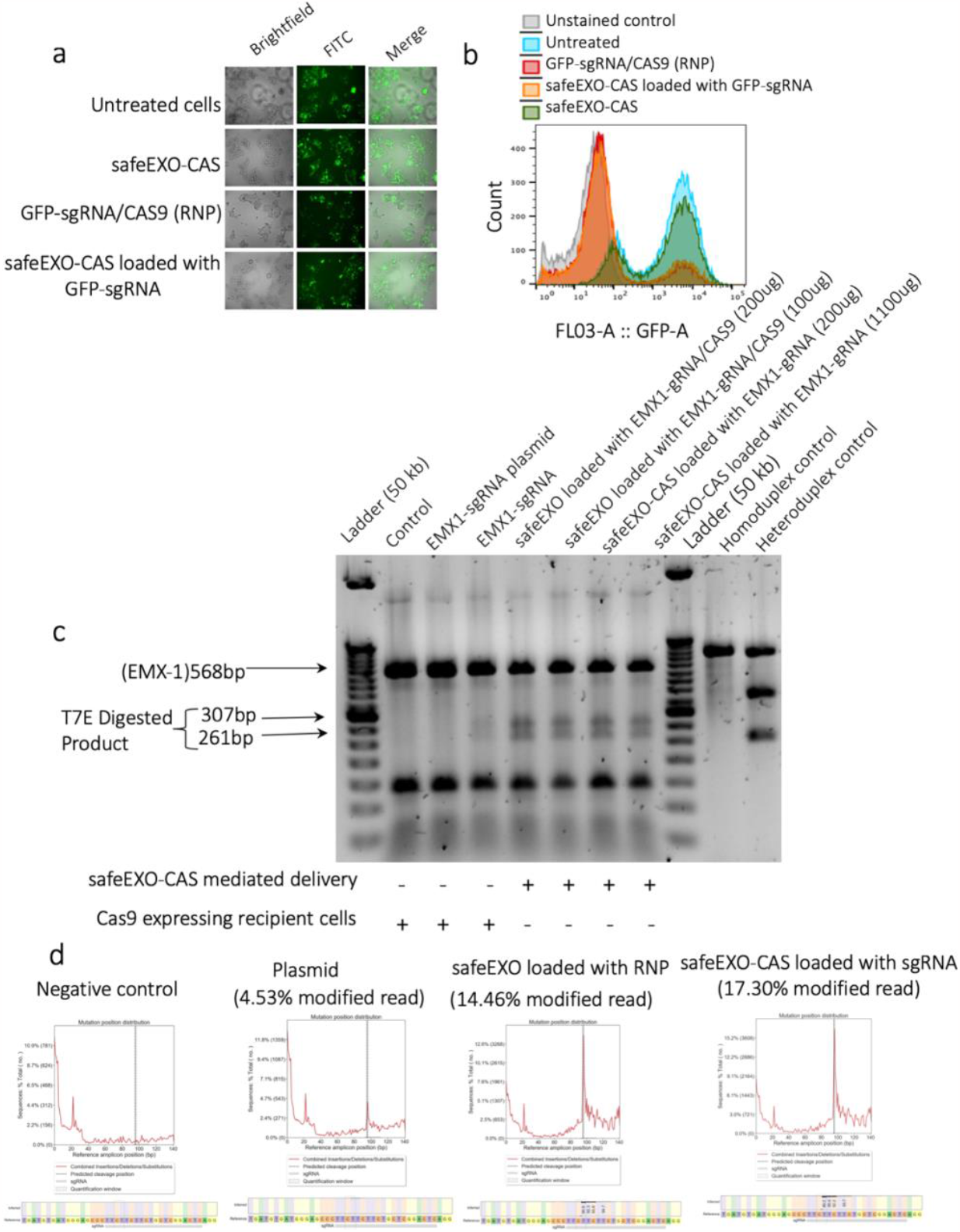
SafeEXO-CAS-mediated non-homologous end joining (NHEJ) genome editing. **a** Fluorescence microscopy and **b** FACS analysis of GFP expressing HEK293T cells treated with safeEXO-CAS, GFP-sgRNA/CAS9 RNP, and safeEXO-CAS loaded with GFP-sgRNA. **c** T7 endonuclease assay against the EMX1 in Cas9 expressing THP-1 cells and in THP-1 cells. Cells were treated with different concentrations of safeEXO or safeEXO-CAS (200ug and 100ug). **d** The percentages of indel induction in EMX1 gene were quantified in each treatment based on deep sequencing of EMX1.

**Fig 5.**
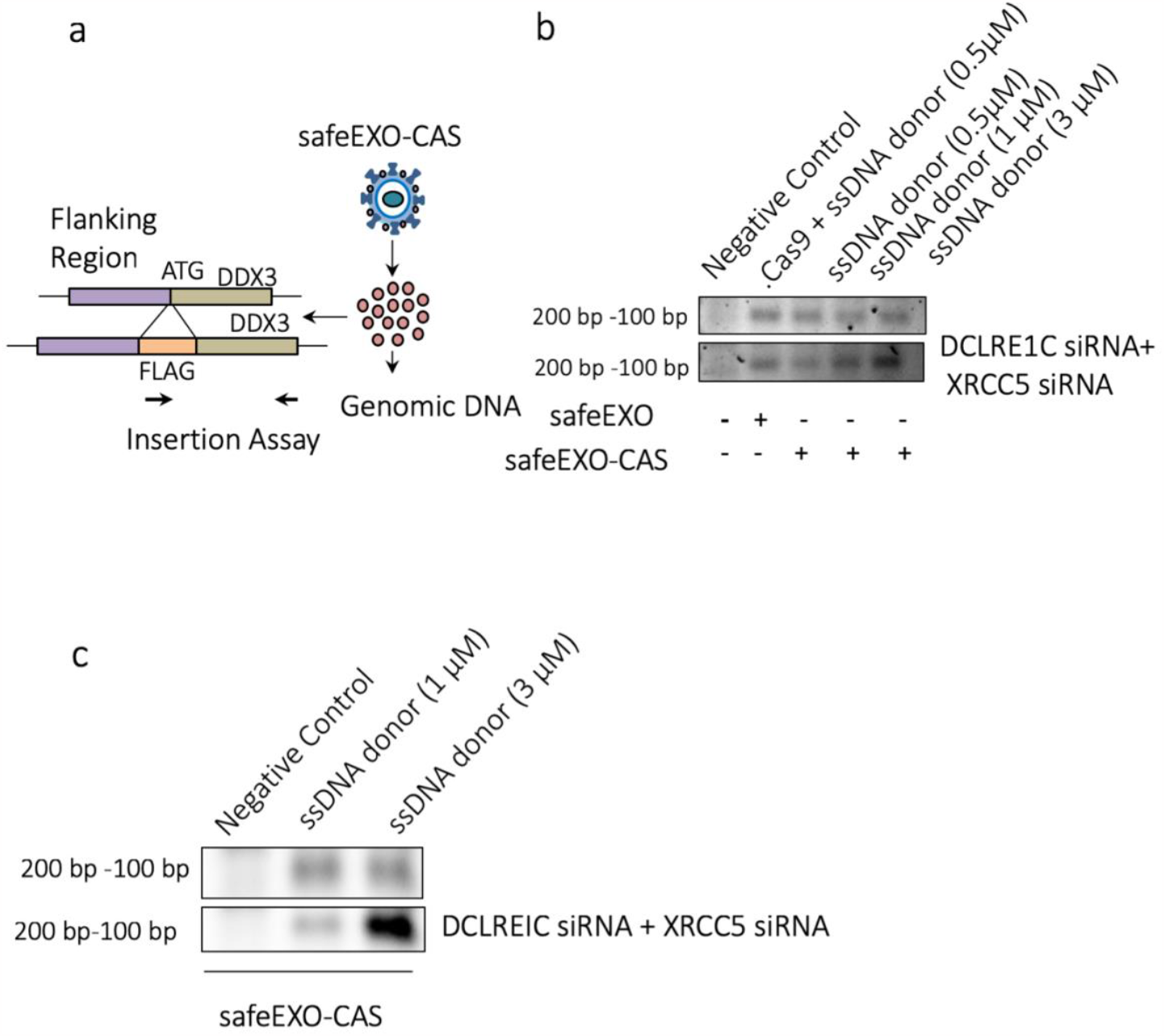
SafeEXO-CAS-mediated homology-directed repair (HDR) editing. **a** Schematic representation of exosome HDR experimental plan in HEK293T cells. **b** SafeEXO-CAS exosomes co-loaded with sgRNA targeting human DDX3 at its start codon and donor ssDNA encoding the FLAG-tag protein surrounded by homology arms to the DDX3 start codon (top). SafeEXO-CAS exosomes were also co-loaded with siRNAs targeting DCLRE1C and XRCC5 to prevent NHEJ (bottom). HEK293T cells were treated with exosomes containing three different concentrations (0.5uM-3uM) of ssDNA donor template. The FLAG-tag insertion was confirmed using the forward primer from FLAG-tag and reverse primer from the DDX3 locus. **c** HEK293T cells were treated with safeEXO-CAS exosomes co-loaded with sgRNA targeting human GFP at its start codon and a ssDNA template encoding the FLAG-tag protein and bearing homology arms to the GFP start codon (top). SafeEXO-CAS exosomes were also co-loaded with siRNAs targeting DCLRE1C and XRCC5 to prevent NHEJ (bottom). HEK293T cells were treated with exosomes containing two different concentrations (0.1uM-3uM) of ssDNA donor template. The FLAG-tag insertion was confirmed using the forward primer from FLAG-tag and reverse primer from GFP locus.

NHEJ genome editing by safeEXO-CAS exosomes was independently confirmed on a different genetic locus in a second cell line. For these confirmatory studies, we used monocytes in suspension culture that are generally refractory to DNA transfection and other gene delivery methods compared to HEK-293 cells. SafeEXO-CAS exosomes were loaded with sgRNA targeting the EMX1 locus and exposed to targeted monocytes in vitro. 72 h after safeEXO-CAS exposure, genome editing at the EMX1 locus was assessed in the bulk cellular population of monocytes using the T7 endonuclease assay and deep-sequencing of the EMX1 locus. Both assays independently demonstrated that safeEXO-CAS exosomes loaded with EXM1-targeted sgRNA efficiently edited the EXM1 locus and was superior to a gene therapy using a plasmid encoding CAS9 and EXM1-targeted sgRNA. Furthermore, safeEXO-CAS exosomes loaded with sgRNA and safeEXO vehicles externally loaded with recombinant Cas9 protein and sgRNA were significantly more efficient compared to the plasmid delivery **(Fig. 4c, 4d)**.

### SafeEXO-CAS vehicles with complex payloads mediate efficient CRISPR-based HDR genomic editing

Precise insertion of genetic material (also known as knock-in) using CRISPR-Cas9 could be accomplished through HDR. HDR is significantly more challenging as a therapeutic modality compared to NHEJ due to decreased efficiency and the added complexity of the payload, which in addition to the Cas9/sgRNA, also must include a donor DNA template. HDR induction was evaluated in target cells after exposure to safeEXO vehicles loaded with the complex payloads needed for HDR, including sgRNA that targeted a locus close to the ATG start codon of the human DDX3 gene along with a template single stranded DNA oligomer encoding the FLAG-tag reporter flanked with 46 nucleotide (nt) homology arms targeting the same region as the sgRNA **(Fig. 5a)**. Cells were exposed to safeEXO-CAS loaded with the sgRNA and ssDNA donor. Cells were passed 5 times after exosome exposure before assessing HDR insertion efficiency by PCR. Cells treated with safeEXO-CAS loaded with sgRNA and ssDNA demonstrated incorporation of the FLAG-tag at the DDX3 locus, proving successful HDR gene editing. A dose dependent increase in the editing efficiency was observed when ssDNA donor concertation was increased from 0.5μM to 3μM **(Fig. 5b)**. To increase HDR efficiency, we co-loaded safeEXO-CAS/sgRNA/ssDNA vehicles with additional siRNAs that target DCLRE1C and XRCC5 genes to prevent non-homologous end joining, resulted in an increase in the insertion of ssDNA to the genome and higher efficiency of HDR **(Fig. 5b)**. HDR genomic editing was independently confirmed in a second cell line and target by exposing HEK-GFP cells with safeEXO-CAS containing sgRNA targeting GFP together with a ssDNA template. HDR was confirmed using a genetic insertion assay (**Fig.5c)**, showing safeEXO-CAS could co-deliver ssDNA, sgRNA, and siRNAs (DCLRE1C and XCRCC5) to efficiently mediate gene insertion in a dose dependent manner **(Fig 5c)**. These data demonstrate the ability of our platform to simultaneously co-deliver highly complex payloads, including CAS, ssDNA, sgRNAs, and siRNAs (DCLRE1C and XCRCC5) to enable efficient HDR-mediated gene insertion, which cannot be efficiently accomplished with other means of delivery.

#### SafeEXO-CAS mediated genomic editing minimizes off-target effect

Initial safety of safeEXO-CAS genome editing was tested looking for cellular inflammatory responses and off target editing. No significant induction of any cytokines was detected after treatment with safeEXO-CAS **(Supplementary Fig.6a)**. Off target editing of safeEXO-CAS was compared to plasmid delivery methods using the standard mismatch tolerance assay utilizing gRNAs with different base mismatches compared to the targeted sequence. Our results demonstrated that safeEXO-CAS genome editing resulted in lower off-target editing compared to the plasmid-based gene editing **(Supplementary Fig.6b)**.

### Targeting safeEXO vehicles to specific biomarkers in vitro and in vivo

To further optimize the safeEXO-CAS exosomes for in vivo therapeutic genomic editing, we tested the potential to engineer them with targeting moieties. SafeEXO-CAS producer cells were engineered to express the integrin alpha chain alpha 6 (ITGA6) on the surface of their exosomes to increase potential lung targeting, as biodistribution studies of circulating exosomes indicated that expression of ITGA6 provides homing to the lung ^30^. To guide the targeting moiety to the exosome surface, producer cells were transfected with cDNA encoding a CD63-ITGA6 protein **(Fig. 6a)**. SafeEXO-CAS-ITGA6 exosomes were collected from these producer cells and flow cytometry quantification of ITGA6 (CD49f) showed significant increase of ITGA6 on the exosome surface **(Fig 6b)**. In a competitive co-culture between targeted lung epithelial cells and non-targeted primary monocytes, safeEXO-CAS-ITGA6 exosomes demonstrated higher uptake in the lung epithelial cells **(Fig. 6c)**.

**Fig 6.**
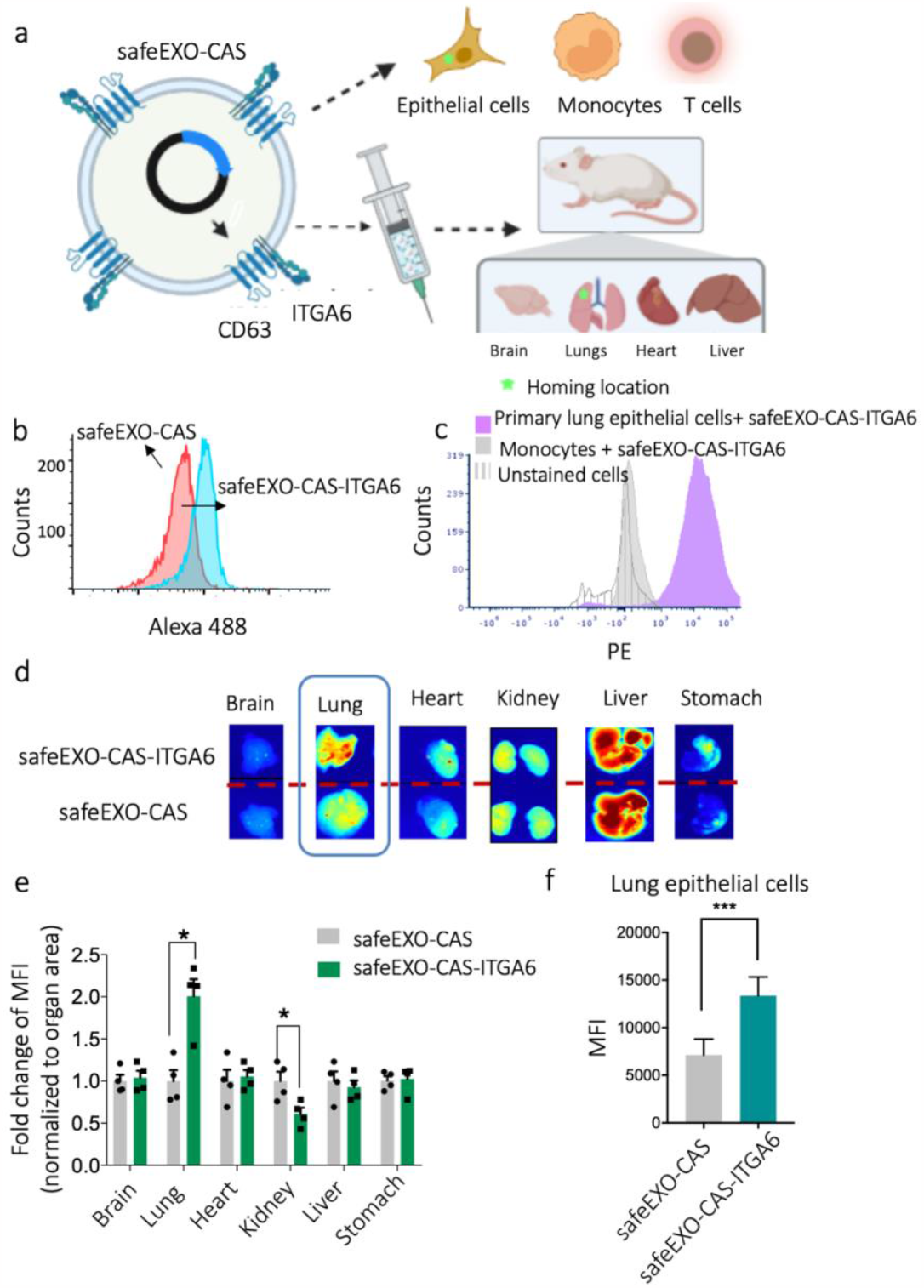
**a** Schematic representation of in vitro and in vivo safeEXO-CAS-ITGA6 lung targeting. **b** Flow cytometry analysis using an ITGA6 (CD49f) antibody of safeEXO-CAS-ITGA6 and untargeted safeEXO-CAS. **c** Flow cytometry analysis of the fluorescently labeled safeEXO-CAS-ITGA6 uptake by primary lung epithelial cells, monocytes and unstained cells after 6h coculture. **d** PKH26 labelled safeEXO-CAS-ITGA6 exosome or untargeted safeEXO-CAS exosome uptake by different mouse organs (brain, lung, heart, kidney, liver and stomach) using optical imaging of organs 15 min after exosome injection. **e** Fold change of fluorescence intensity normalized to organ area from different organs (n=8 total). **f** Percentage of exosome uptake was quantified by flow cytometry in lung epithelial cells (EpCAM^+^) of mice receiving safeEXO-CAS-ITGA6 or untargeted safeEXO-CAS control exosomes (n=6 per group). Data are represented as mean ± standard deviation (SD). MFI, mean fluorescent intensity, \**p* ≤ 0.05, Student’s T test was used for pairwise comparison.

To test in vivo targeting, biodistribution studies were performed on healthy mice after injection of NIR-labeled safeEXO-CAS-ITGA6 exosomes versus control NIR-labeled untargeted safeEXO-CAS exosomes. SafeEXO-CAS-ITGA6 exosomes demonstrated higher uptake in the lung **(Fig 6 d, e)** and corresponding lower uptake in kidney compared to control untargeted safeEXO-CAS exosomes. Flow cytometry of lung epithelial cells (EpCAM^+^) showed an increased presence of exosomes in the lung epithelial cells isolated from mice which received safeEXO-CAS-ITGA6 exosomes compared to the controls **(Fig. 6f)**. These data indicate that expression of ITGA6 on surface of safeEXO-CAS exosomes increases their uptake in lung epithelial cells. The liver demonstrated intense uptake of both targeted and untargeted exosomes, which has been shown by many groups, as the liver plays a primary role in clearing extracellular vesicles from circulation. However, only the specific binding to the lung by targeted safeEXO-CAS-ITGA6 exosomes demonstrated successful genomic editing, in contrast to the liver.

### SafeEXO-CAS-ITGA6 can mediated successful genome editing in lung

Genomic editing was tested by loading safeEXO-CAS-ITGA6 exosomes with 3 sgRNAs designed to disrupt the reading frame of EXM1 through indels caused by NHEJ. sgRNA-loaded safeEXO-CAS-ITGA6 exosomes or control vehicle (unloaded safeEXO-CAS) were systemically injected into mice, which were analyzed for genome editing two weeks after injection **(Fig. 7a)**. All mice injected with safeEXO-CAS-ITGA6 loaded with sgRNA targeting EMX1 displayed significant editing in the lung (between 7% and 16% efficiency; **Fig. 7b**) compared to the no editing detected in lungs of control mice. In contrast, no significant editing was seen in other organs, including in the liver **(Fig. 7c)**, despite the high localization of the exosomes in the biodistribution studies (**Fig. 6d**). These data indicate that the untargeted liver uptake is likely due to the exosome clearance function of the liver and does not indicate functional targeting.

**Fig 7.**
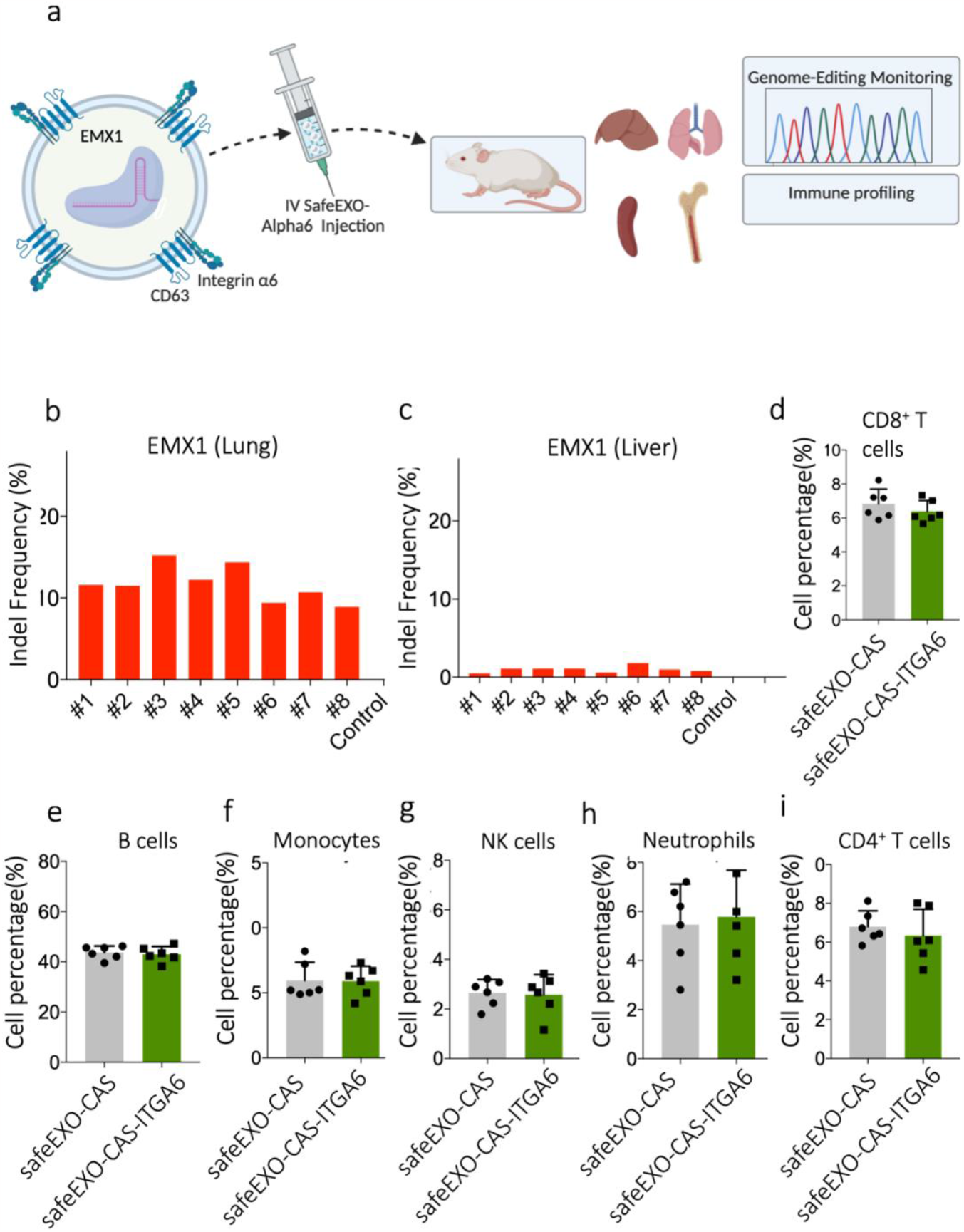
**a** Schematic representation of safeEXO-CAS-ITGA6 mediated EMX1 editing in vivo. **b** Percentage of indel frequency in EMX1 in lung and **c** liver (n=8 per group). **d-i** Percentage of immune subtypes frequency in CD8^+^T cells, B cells, monocytes, NK cells, neutrophils, and CD^+^4 T cells (n=6 per group). Data are represented as mean ± standard deviation (SD).

### Initial toxicology studies after safeEXO mediated genomic editing

To assess for adverse effects of our safeEXO-CAS-ITGA6 genomic edition vehicle, we observed for clinical morbidity, and performed immunoprofiling and plasma toxicology studies on the EXM1-edited mice. There were no associated signs of morbidity and immuneprofiling of safeEXO-CAS-ITGA6 treated mice and control mice did not show any detectable changes in the frequencies of immune cells (CD8 T cells, CD4 T cells, NK cells, monocytes, B cells and neutrophils) in the spleen **(Fig 7d-i)** and bone marrow **(Supplementary Fig.7)**. Blood biochemical and toxicological panels did not reveal any systemic adverse effects of safeEXO-CAS-ITGA6 exosomes (**Supplementary Fig.8)**.

## DISCUSSION

The results presented in this study demonstrate the potential of the safeEXO platform as a novel, non-invasive method to deliver RNAi and CRISPR-based gene editing components for high efficiency delivery and genomic editing. We showed that safeEXO-CAS exosomes can effectively deliver highly complex payloads, including sgRNA, ssDNA and siRNAs, a novel strategy that we demonstrate can block NHEJ and facilitate HDR-based gene insertions, which is not feasible with other delivery modalities. Furthermore, we engineered the safeEXO-CAS exosomes to express ITGA6 on their surface as an example of tissue/biomarker targeting and showed their potential to facilitate efficient in vivo gene editing in mouse lung at 10-15% efficiency after a single injection.

Our findings suggest that safeEXO-CAS exosomes have a wide range of potential applications, from in vitro to preclinical studies, and potentially future clinical exploration. safeEXO-CAS exosomes could be used to study gene functions and develop novel therapies for different diseases ranging from monogenetic disorders to cancer. As safeEXO-CAS exosomes were non-immunogenic, they have the potential to be used in vivo to target specific organs or tissues with CRISPR components without eliciting an immune response. For example, our prototype targeted safeEXO-CAS exosomes displaying integrin alpha-6 (ITGA6) on their surface could be useful for the treatment of lung diseases, as our results demonstrate they can deliver functional CRISPR components to the lungs and mediate targeted efficient genome editing preferentially to targeted tissue and not the liver.

The successful generation of exosome-based drug delivery platforms for the targeted delivery of CRISPR/Cas offers numerous advantages over existing viral or non-viral vectors. These engineered exosomes are biocompatible, non-immunogenic, and non-toxic, even in repeated in vivo injections. Furthermore, their lipid bilayer protects the protein and RNA cargos from enzymes such as proteases and RNases, allowing for long circulation times. Finally, exosomes carrying bacterial-derived Cas machinery have been demonstrated to exhibit an increased capacity to escape degradation or clearance by the immune system compared with existing CRISPR delivery vectors. The successful delivery of functional CRISPR/Cas systems using exosomes, as demonstrated here, provides a promising and powerful platform for gene editing in vitro and in vivo.

This work provides a promising step towards achieving effective, targeted delivery of CRISPR/Cas systems for gene editing in a scalable and safe manner. Further research is needed to identify the optimal combination of nucleic acid and Cas9 components, and exosome dosing in the context of disease. Taken together, this work provides an exciting platform for future development of exosome-based gene editing applications. If successful, the safeEXO-CAS platform could impact precision medicine by providing an effective and scalable gene editing therapy with minimal off-target effects. This would enable personalized therapeutics for a wide range of diseases caused by genetic mutations, leading to more targeted treatments that improve patient outcomes.

## METHODS

### Cell culture and transfections

Cas9 expressing THP-1 monocytes, THP-1, and KRAS 4B wild type cells were cultured in RPMI-1640 and HEK293T in DMEM, supplemented with 10% Fetal Bovine Serum (FBS) (Corning #35-011-CV) containing 1X Penicillin and Streptomycin mix (100X penicillin (10,000IU) and streptomycin (10,000μg/ml) mix) (Corning #30-002-CI). All cell lines were cultured and incubated at 37 °C with 5% CO2. Cas9 expressing THP-1 monocytes were transfected with fluorescently labeled sgRNA against Drosha, ALIX, and hnRNPA2B1. cells were sorted and individual knockout clones’ selection was confirmed using western blot.

All experiments involving exosome isolation were performed in media supplemented with exosome depleted fetal bovine serum (Gibco #A27208-03). For generation of Drosha, Alix, and hnRNPA2B1 knockout cells, cells transfected with 2sgRNAs against each gene and screening by western blot as described below to identify the knockout clones.

### Collection and purification of exosomes

In a T75 or T25 flasks producer cells of exosomes were seeded and 24 h later transfected with 24 μg total of plasmids or vectors if indicated. After 12 h of transfection, the cell culture medium was replaced. After 72 h of transfection, the media was collected and subjected to sequential spins: a low-level spin at 800 × *g* for 5 mins to remove cells, followed by 2000 × *g* for 20 mins to remove of cell debris. ExoQuick-TC ULTRA (for cell culture medium) (System Biosciences) was used according to the manufacturer’s protocol. The ExoQuick was added followed by an incubation overnight at 4°C following the protocol recommended for the manufacturer. After incubation, the precipitate was spun down (3000 × *g*, 10 min at 4°C), exosomes were resuspended in buffer and added to the purification columns and exosomes were eluted in volume of 500 μl elution buffer per isolation. The final volume was passed through a 0.22-micron filter (Millipore Sigma). The exosome isolated fraction and EV-depleted fraction were frozen and stored at -80°C until further use.

### Exosome surface marker analysis

The exosomes were incubated with capture beads for CD81 (ab239687, ABCAM) and CD63 (ab239686, ABCAM) overnight at 4 °C, as recommended by the manufacturer. Primary detection antibody mixture (CD49f, CD63 and CD81 conjugated to fluorescent dyes) was added to the mixture of samples and beads (1:25). Samples were mixed gently and incubated for 1h at 4°C. Samples were washed 2 times by assay buffer and resuspended in 350ul FACS buffer. Samples were run on a BD FACSAria II and data was analyzed using FCS Express Analysis Software (De Novo Software, Pasadena, CA).

### Protein quantification and immunoblot analysis

Exosomes along with whole cell pellets were lysed in RIPA Lysis and Extraction Buffer (Thermo Fisher #89900). Protein concentration was determined by a Pierce BCA Protein Assay Kit (Thermo Fisher #23227) as per manufacturer’s instructions.

Exosomes and cell lysates were lysed in RIPA buffer and run on a 15% polyacrylamide gel with equal amounts of protein loaded (100 μg). Proteins were transferred to nitrocellulose membrane (Bio-Rad #1620115) and then blocked for 1 hours in 1X TBS 1% Casein Blocker (blocking buffer) (Bio-Rad #1610782).

The following primary antibodies were used: Exosome-anti-CD81 antibody (Invitrogen #10630D), Exosome-anti-CD63 antibody (Invitrogen #10628D), Drosha Antibody (Invitrogen #PA5-79927), hnRPA2B1 Polyclonal Antibody (Invitrogen #PA5-34939), Cas9 Antibody (10C11-A12) (Invitrogen #MA1-202), Alix Antibody (3A9) (Invitrogen #MA1-83977). All primary antibodies were used at a dilution of 1:1000 in blocking buffer and were incubated overnight. For detection, secondary goat anti-mouse IgG (H+L)-HRP Conjugate antibody (Bio-Rad #170-6516) and goat anti-rabbit IgG (H+L)-HRP Conjugate antibody (Bio-Rad #170-6515) was used for 2 hours at a dilution rate of 1:3000 in blocking buffer. The immunoreactive bands were visualized by a Clarity Max™ Western ECL substrate (Bio-Rad #1705062) according to the manufacturer’s protocol and an iBright Imaging systems (Thermo Fisher Scientific #CL1000).

### Mice and in vivo biodistribution studies

C57BL/6J mice (JAX stock #000664, Jackson Laboratory) (6-8 weeks old) were maintained in accordance with the Guide for the Care and Use of Laboratory Animals. Animal housing follows a 12 h light/12 h dark cycle, wherein the temperature is maintained between 68–79 °F. Humidity was maintained between 30–70 percent. All experiments were performed according to the guidelines of the Institutional Animal Committee of the Columbia University.

Exosomes were labeled with ExoGlow (EXOGM600A-1, SBI) as described by the manufacturer. Briefly, 1μl of ExoGlow was added to 100μg of exosomes in 200μl 1X PBS and incubated at room temperature for 1hr. Labeled exosomes were mixed with 63μl of ExoQuick-TC and incubated overnight at 4°C followed by centrifugation at 13,000 *g* for 10 mins. The resulting pellet containing labeled exosomes were resuspended in 1X PBS buffer. Further, 100 μg of exosomes were administered into mice in a total of 100 μl volume via IV injection. 15-30 minutes after exosome injection, mice were euthanized using an overdose of isoflurane anesthesia. The animals and organs (brain, lung, heart liver, and spleen) were imaged using an IVIS spectrum bioimaging device (PerkinElmer). The intensity of the fluorescence was calculated using the ImageJ and normalized to the controls. Blood was collected for toxicological and biological assessments.

### Nanoparticle tracking analysis (NTA)

The concentration and size of exosomes in cultured media were identified by a NanoSight NS300 system (NanoSight, Amesbury, UK) supplied with a fast video capture and Nanoparticle Tracking Analysis (NTA) software. Before performing the experiments, the instrument was calibrated with 100 nm polystyrene beads (Thermo Scientific, Fremont, CA, USA). The samples were imaged three times each for 30s at 25°C. NTA software processed the video captures and measured the associated particle concentrations (particles/ml), size distributions (in nanometer), and intensities (arb. units) of the samples.

### sgRNA design and sequences (+PAM)

sgRNAs targeting *DDX3, GFP, EMX1* were designed using IDT software. Human *DDX3*: 5′ AGGGATGAGTCATGTGGCAGtgg 3′ Human *EMX1*: 5′ GAGTCCGAGCAGAAGAAGAAggg 3′ Mouse *EMX1_SG1*: 5′ CAAGCGACGTTCCCCAGGACggg 3′ Mouse *EMX1_SG2*: 5′ CCAAGGATGGTGGCACCGGCggg 3′ Mouse *EMX1_SG3*: 5′ GGCAGGGAAGCCACTCACGAagg 3′ *GFP*: 5′ CGGCCATGATATAGACGTTGtgg 3′

### T7 endonuclease assay

Genomic DNA was extracted from cells using the Quick-DNA Miniprep Kit (Zymo Research # D3024). 40ng of genomic DNA was then used for PCR amplification for 25μl reaction. Further, in order to form heteroduplexes Alt-R Genome Editing Detection Kit (IDT # 1075932) was used. Briefly, 10μl PCR products were added with T7EI reaction buffer (10X) and 6μl H20 to a final volume of 18μl. The reaction was heated and cooled according to the following procedure: heated at 95°C for 10 min, ramping from 95-85°C at a ramp rate of -2°C/sec (ramp1), then ramping from 85-25°C at a ramp rate of -0.3°C/sec (ramp2). For T7EI digestion, 2μl of T7 endonuclease I(1U/μl) was added to the heteroduplexes. The reaction was incubated at 37°C for 60 minutes. Samples were finally run on a 2.5% agarose gel. For T7 endonuclease assay the primers used were: GFP_F: GCAAGGGCGAGGAGCTGTTCAC GFP_R: AGGTAGTGGTTGTCGGGCAGCAG EMX1_T7endoF: TTCTCTCTGGCCCACTGTGTCCTC EMX1_T7endoR: AGCCCATTGCTTGTCCCTCTGTCAATG

### Exosome mediated homology directed repair using ssDNA

Exosomes were loaded with different concentration of ssDNA donor template containing FLAG-tag (5′-ACTCGCTTAGCAGCGGAAGACTCCGagTTCTCGGTACTCTTCAGGGATGGA CTACAAGGACGACGATGACAAGagTCATGTGGCAGTGGAAAATGCGCTCGGGCTGG ACCAGCAGGTGA-3’) targeting AUG codon of DDX3 at 0.5 μM, 1 μM and 3 μM with or without addition of DCLRE1C siRNA (cat# AM16708) and XRCC5 siRNA (cat# AM16708). 8X10^4^ THP-1 (0.5 μM, 1 μM and 3 μM of ssDNA) or HEK293T (1 μM and 3 μM of ssDNA) cells were treated with exosomes. cells were then collected for the genomic DNA isolation and PCR. Cells were checked for the insertion using the following primers: FLAG-tag-Forward 5′-GACTACAAGGACGACGATGACAAG-3′ and DDX3-Reverse2 5′-CGCCATTAGCCAGGTTAGGT-3′.

### High-throughput sequencing of Emx1

Genomic DNA was extracted from exosome-treated cells using the Genomic DNA extraction kit (Zymo Research, Irvine, CA). 40 ng of genomic DNA was then used for PCR amplification using primers specific for EMX1 (EMX1-Forward 5′-GGCCCAGGTGAAGGTGTGGTT -3′ and EMX1-Reverse 5′-GGTTGCCCACCCTAGTCATTGGA -3′). Obtained PCR products were gel purified and sequenced at the MGH Center for Computational and Integrative Biology (CCIB) at Boston, Massachusetts. The CRISPR editing efficiency was identified using CRISPResso2^22^.

### IV injection of safeEXO-CAS

Genomic DNA from each mouse (treated either by control or lung targeting safeEXO-CAS) was extracted from lung and liver. Following this, a PCR was performed on 40 ng of gDNA template, the PCR was performed using the following primers: EMX1_sgRNA2_Forward: TTAGGGCTCTCGCACGCCCCTC, EMX1_sgRNA2_Reverse: TGGTTCATGGCCTCTGGGAACACCA, EMX1_sgRNA3_Forward: TGCACACCCCGCACGGCGGCA EMX1_sgRNA3_Reverse:\ CCTGGAAGCGGTGGCCAAAGAAGCGA, to amplify the EMX1 amplicon (95 °C -5 min, 35 cycles (95 °C -30 sec, 60 °C -30sec, 72 °C -45sec), 72 °C -5 min). Amplicons were run on 1% Agarose gel and purified after excision. Further, PCR amplified products were analyzed by sanger sequencing (GENEWIZ, NJ) electropherograms and TIDE analysis.

### Generation of single-cell suspensions and flow cytometry analysis

Single-cell suspensions of the spleen and bone marrow were prepared for analysis. Spleens were homogenized through a 70 μm cell strainer, while bone marrow was flushed into a falcon tube with a syringe. Erythrocytes in the suspension were lysed by ACK lysing buffer for 4 minutes, and the reaction was then stopped with PBS 0.1% BSA. Finally, the suspension was passed through a 30 μm cell strainer. Lung tissue was dissociated using a Miltenyi dissociation kit. The lung epithelial cells were harvested by positive selection using EpCAM (cat#130-105-958, Miltenyi), the pan-epithelial marker based on the manufacturer’s protocol. Flow cytometry for identification of cell counts and mean fluorescent intensity of immune cells was performed on a BD FACSAria II. Immune cell populations were defined on live lymphocytes with CD45 and Zombie Aqua™ (cat# 423101) and CD4^+^ T cells (TCRβ^+^CD4^+^), CD8^+^ T cells (CD4^-^CD8^+^), NK (TCRβ^-^NK1.1^+^), neutrophils (Ly6G^+^CD11b^+^), monocytes (TCRβ^-^Ly6G^-^CD11b^+^Ly6C^-^), B cells (TCRβ^-^CD19^+^). Flow cytometry data was analyzed using FCS Express Analysis Software (De Novo Software, Pasadena, CA). The following antibody clones were used for this study: CD45 (Alexa Fluor 488, 30-F11, BioLegend), TCRβ (BUV737, H57-597, BD), CD8a (Alexa 700, 53-6.7, BioLegend), CD4 (BV605, RM4-5, BioLegend), NK-1.1 (Brilliant Violet 650, PK136, BioLegend), Ly6G (BUV395, 1A8, BD), Ly6C (BV786, HK1.4, BioLegend), CD11b (BV421, M1/70, BioLegend), CD19^+^(PE, 4G7, BioLegend).

### Statistical analysis

Data presented as mean+ standard deviation (SD) . Data analyses was performed by GraphPad Prism 6.01 (GraphPad, USA) and R software (V 3.6). Differences between two groups were tested using the Student’s *t* test. Differences among multiple groups were analyzed using one-way ANOVA followed by Dunnett’s post hos comparisons. *P* value <0.05 was considered significant. Experiments were repeated at least two times with at least three replicates. Statistical significance was annotated as follows: *p < 0.05, **p < 0.01, ***p < 0.001, ***p<0.0001.

## References

1 Momen-Heravi, F. et al. Current methods for the isolation of extracellular vesicles. Biol Chem 394, 1253–1262, doi:10.1515/hsz-2013-0141 (2013).

2 Momen-Heravi, F., Getting, S. J. & Moschos, S. A. Extracellular vesicles and their nucleic acids for biomarker discovery. Pharmacol Ther 192, 170–187, doi:10.1016/j.pharmthera.2018.08.002 (2018).

3 Lotvall, J. et al. Minimal experimental requirements for definition of extracellular vesicles and their functions: a position statement from the International Society for Extracellular Vesicles. J Extracell Vesicles 3, 26913, doi:10.3402/jev.v3.26913 (2014).

4 Thery, C., Zitvogel, L. & Amigorena, S. Exosomes: composition, biogenesis and function. Nat Rev Immunol 2, 569–579, doi:10.1038/nri855 (2002).

5 Momen-Heravi, F., Bala, S., Bukong, T. & Szabo, G. Exosome-mediated delivery of functionally active miRNA-155 inhibitor to macrophages. Nanomedicine 10, 1517–1527, doi:10.1016/j.nano.2014.03.014 (2014).

6 Malhotra, H. et al. Exosomes: Tunable Nano Vehicles for Macromolecular Delivery of Transferrin and Lactoferrin to Specific Intracellular Compartment. J Biomed Nanotechnol 12, 1101–1114, doi:10.1166/jbn.2016.2229 (2016).

7 Hood, J. L. Post isolation modification of exosomes for nanomedicine applications. Nanomedicine (Lond) 11, 1745–1756, doi:10.2217/nnm-2016-0102 (2016).

8 Ishida, T., Harashima, H. & Kiwada, H. Liposome clearance. Biosci Rep 22, 197–224, doi:10.1023/a:1020134521778 (2002).

9 Bukong, T. N., Momen-Heravi, F., Kodys, K., Bala, S. & Szabo, G. Exosomes from hepatitis C infected patients transmit HCV infection and contain replication competent viral RNA in complex with Ago2-miR122-HSP90. PLoS Pathog 10, e1004424, doi:10.1371/journal.ppat.1004424 (2014).

10 Bala, S. et al. Biodistribution and function of extracellular miRNA-155 in mice. Sci Rep 5, 10721, doi:10.1038/srep10721 (2015).

11 Salsman, J., Masson, J. Y., Orthwein, A. & Dellaire, G. CRISPR/Cas9 Gene Editing: From Basic Mechanisms to Improved Strategies for Enhanced Genome Engineering In Vivo. Curr Gene Ther 17, 263–274, doi:10.2174/1566523217666171122094629 (2017).

12 Hentze, M. W. & Kulozik, A. E. A perfect message: RNA surveillance and nonsense-mediated decay. Cell 96, 307–310, doi:10.1016/s0092-8674(00)80542-5 (1999).

13 Miller, J. C. et al. A TALE nuclease architecture for efficient genome editing. Nat Biotechnol 29, 143–148, doi:10.1038/nbt.1755 (2011).

14 Urnov, F. D. et al. Highly efficient endogenous human gene correction using designed zinc-finger nucleases. Nature 435, 646–651, doi:10.1038/nature03556 (2005).

15 Lino, C. A., Harper, J. C., Carney, J. P. & Timlin, J. A. Delivering CRISPR: a review of the challenges and approaches. Drug Deliv 25, 1234–1257, doi:10.1080/10717544.2018.1474964 (2018).

16 Zhang, S., Shen, J., Li, D. & Cheng, Y. Strategies in the delivery of Cas9 ribonucleoprotein for CRISPR/Cas9 genome editing. Theranostics 11, 614–648, doi:10.7150/thno.47007 (2021).

17 Wang, A. Y., Peng, P. D., Ehrhardt, A., Storm, T. A. & Kay, M. A. Comparison of adenoviral and adeno-associated viral vectors for pancreatic gene delivery in vivo. Hum Gene Ther 15, 405–413, doi:10.1089/104303404322959551 (2004).

18 Banaszynski, L. A., Chen, L. C., Maynard-Smith, L. A., Ooi, A. G. & Wandless, T. J. A rapid, reversible, and tunable method to regulate protein function in living cells using synthetic small molecules. Cell 126, 995–1004, doi:10.1016/j.cell.2006.07.025 (2006).

19 Wilbie, D., Walther, J. & Mastrobattista, E. Delivery Aspects of CRISPR/Cas for in Vivo Genome Editing. Acc Chem Res 52, 1555–1564, doi:10.1021/acs.accounts.9b00106 (2019).

20 Peer, D. et al. Nanocarriers as an emerging platform for cancer therapy. Nat Nanotechnol 2, 751–760, doi:10.1038/nnano.2007.387 (2007).

21 Mout, R., Ray, M., Lee, Y. W., Scaletti, F. & Rotello, V. M. In Vivo Delivery of CRISPR/Cas9 for Therapeutic Gene Editing: Progress and Challenges. Bioconjug Chem 28, 880–884, doi:10.1021/acs.bioconjchem.7b00057 (2017).

22 Clement, K. et al. CRISPResso2 provides accurate and rapid genome editing sequence analysis. Nat Biotechnol 37, 224–226, doi:10.1038/s41587-019-0032-3 (2019).

23 Momen-Heravi, F., Bala, S., Kodys, K. & Szabo, G. Exosomes derived from alcohol-treated hepatocytes horizontally transfer liver specific miRNA-122 and sensitize monocytes to LPS. Sci Rep 5, 9991, doi:10.1038/srep09991 (2015).

24 Warnecke-Eberz, U., Chon, S. H., Holscher, A. H., Drebber, U. & Bollschweiler, E. Exosomal onco-miRs from serum of patients with adenocarcinoma of the esophagus: comparison of miRNA profiles of exosomes and matching tumor. Tumour Biol 36, 4643–4653, doi:10.1007/s13277-015-3112-0 (2015).

25 Geis-Asteggiante, L. et al. Differential Content of Proteins, mRNAs, and miRNAs Suggests that MDSC and Their Exosomes May Mediate Distinct Immune Suppressive Functions. J Proteome Res 17, 486–498, doi:10.1021/acs.jproteome.7b00646 (2018).

26 Villarroya-Beltri, C. et al. Sumoylated hnRNPA2B1 controls the sorting of miRNAs into exosomes through binding to specific motifs. Nat Commun 4, 2980, doi:10.1038/ncomms3980 (2013).

27 Schwarzenbach, H. & Gahan, P. B. MicroRNA Shuttle from Cell-To-Cell by Exosomes and Its Impact in Cancer. Noncoding RNA 5, doi:10.3390/ncrna5010028 (2019).

28 Iavello, A. et al. Role of Alix in miRNA packaging during extracellular vesicle biogenesis. Int J Mol Med 37, 958–966, doi:10.3892/ijmm.2016.2488 (2016).

29 Pucihar, G., Kotnik, T. & Miklavcic, D. Measuring the induced membrane voltage with Di-8-ANEPPS. J Vis Exp, doi:10.3791/1659 (2009).

30 Hoshino, A. et al. Tumour exosome integrins determine organotropic metastasis. Nature 527, 329–335, doi:10.1038/nature15756 (2015).

